# Stable recurrent dynamics in heterogeneous neuromorphic computing systems using excitatory and inhibitory plasticity

**DOI:** 10.1101/2023.08.14.553298

**Authors:** Maryada, Saray Soldado-Magraner, Martino Sorbaro, Rodrigo Laje, Dean V. Buonomano, Giacomo Indiveri

**Affiliations:** Institute of Neuroinformatics, University of Zurich and ETH Zurich, Zurich, Switzerland; Department of Neurobiology, University of California, Los Angeles, USA; ETH AI Center, ETH Zurich, Switzerland; Department of Science and Technology, Universidad Nacional de Quilmes, Bernal, Argentina; CONICET, Argentina

## Abstract

Many neural computations emerge from self-sustained patterns of activity in recurrent neural circuits, which rely on balanced excitation and inhibition. Neuromorphic electronic circuits that use the physics of silicon to emulate neuronal dynamics represent a promising approach for implementing the brain’s computational primitives, including self-sustained neural activity. However, achieving the same robustness of biological networks in neuromorphic computing systems remains a challenge, due to the high degree of heterogeneity and variability of their analog components.

Inspired by the strategies used by real cortical networks, we apply a biologically-plausible cross-homeostatic learning rule to balance excitation and inhibition in neuromorphic implementations of spiking recurrent neural networks. We demonstrate how this learning rule allows the neuromorphic system to work in the presence of device mismatch and to autonomously tune the spiking network to produce robust, self-sustained, fixed-point attractor dynamics with irregular spiking in an inhibition-stabilized regime. We show that this rule can implement multiple, coexisting stable memories, with emergent soft-winner-take-all (sWTA) dynamics, and reproduce the so-called “paradoxical effect” widely observed in cortical circuits. In addition to validating neuroscience models on a substrate that shares many similar properties and limitations with biological systems, this work enables the construction of ultra-low power, mixed-signal neuromorphic technologies that can be automatically configured to compute reliably, despite the large on-chip and chip-to-chip variability of their analog components.

## Introduction

Animal brains can perform complex computations including sensory processing, motor control, and working memory, as well as higher cognitive functions, such as decision-making and reasoning, in an efficient and reliable manner. At the neural network level, these processes are implemented using a variety of computational primitives that rely on neural dynamics within recurrent neocortical microcircuits [1–6]. Translating the computational primitives observed in the brain into novel technologies can potentially lead to radical innovations in artificial intelligence and edge-computing applications. A promising technology that can implement these primitives with compact and low-power devices is that of mixed-signal neuromorphic systems, which employ analog electronic circuits to emulate the biophysics of real neurons [7–9]. As opposed to software simulations, the direct emulation performed by these electronic circuits relies on the use of the physics of the silicon substrate to faithfully reproduce the dynamics of neurons and synapses asynchronously in real time. While systems designed following this approach have the advantage of ultra-low power consumption, they have limitations and constraints similar to those found in biology. These include a high degree of variability, heterogeneity, and sensitivity to noise [10, 11]. Due to these constraints, implementing recurrent networks in silicon with the same stability and robustness observed in biological circuits has remained an open challenge.

Previous studies have already demonstrated neuromorphic implementations of recurrent computational primitives such as winner-take-all or state-dependent networks [12–15]. However, these systems required manual tuning and exhibited dynamics that would not always produce stable self-sustained regimes. Manually tuning on-chip recurrent networks with large numbers of parameters is challenging, tedious, and not scalable. Furthermore, because of cross-chip variability, this process requires re-tuning for each individual chip. Even when a stable regime is achieved with manual tuning, changes in the network conditions (e.g., due to temperature variations, increase or decrease of input signals, etc.) would require re-tuning. Therefore, mixed-signal neuromorphic technology can greatly benefit from automatic stabilizing and tuning mechanisms. While not aiming to remove existing heterogeneities (which are also present in biological systems), we can work with them to manage the challenges imposed by the analog substrate.

Neocortical computations rely heavily on positive feedback imposed by recurrent connections between excitatory neurons, which allows networks to perform complex time-dependent computations and actively maintain information about past events. Recurrent excitation, however, also makes neocortical circuits vulnerable to “runaway excitation” and epileptic activity [16, 17]. In order to harness the computational power of recurrent excitation and avoid pathological regimes, the neocortical circuits operate in an inhibition-stabilized regime, in which positive feedback is held in check by recurrent inhibition [18–21]. Indeed, there is evidence that the default awake cortical dynamic regime may be inhibition-stabilized [20, 22]. It is generally accepted that neocortical microcircuits have synaptic learning rules in place to homeostatically balance excitation and inhibition in order to generate dynamic regimes capable of self-sustained activity [23–27]. Recently, a family of synaptic learning rules that differentially operate at excitatory and inhibitory synapses has been proposed, which can drive simulated neural networks to self-sustained and inhibition-stabilized regimes in a self-organizing manner [28]. This family of learning rules is referred to as being “cross-homeostatic” because synaptic plasticity at excitatory and inhibitory synapses is dependent on both the inhibitory and excitatory setpoints.

Taking inspiration from self-calibration principles in cortical networks, we use these cross-homeostatic plasticity rules to guide neuromorphic circuits to balanced excitatory-inhibitory regimes in a self-organizing manner.

We show that these rules can be successfully employed to autonomously calibrate analog spiking recurrent networks in silicon. Specifically, by automatically tuning all synaptic weight classes in parallel, the dynamics of the silicon networks converge to a fully self-sustained inhibition-stabilized regime. The weights and firing rates converge to stable, fixed-point attractors. In addition, the emergent (fast) neural dynamics reaches an asynchronous, irregular regime and express the “paradoxical effect”, a signature of inhibition-stabilized networks widely observed in cortical circuits, where an increase in input to the inhibitory population results in a counter-intuitive decrease in its firing response [18, 20]. The plasticity rules proposed prove resilient to hardware variability and noise, also across different chips, as well as to different parameter initializations. Importantly, we demonstrate that inhibitory plasticity (often neglected in neuromorphic electronic systems) is necessary for successful convergence. We also demonstrate that by utilizing these plasticity rules, multiple, coexisting long-term memories can be maintained within the same network without disrupting the learned dynamics. Additionally, we show that several such networks can be robustly implemented on a single chip. Interestingly, the different memory ensembles show emergent sWTA dynamics, even though no specific inhibitory competition is externally imposed into the network.

From a computational neuroscience perspective, these results validate the robustness of cross-homeostatic plasticity in a physical substrate that presents similar challenges to those of biological networks, unlike idealized digital simulations. From a neuromorphic perspective, this approach provides the community with a reliable method to autonomously and robustly calibrate recurrent neural networks in future mixed-signal analog/digital systems that are affected by device variability (also including memristive devices [29, 30]). This will open the door to designing autonomous systems that can interact in real time with the environment, and compute reliably with low-latency at extremely low-power using attractor dynamics, as observed in biological circuits.

## Results

In the cortex, excitatory Pyramidal (Pyr) and inhibitory Parvalbumin-positive interneurons (PV) constitute the main neuronal subtypes and are primarily responsible for excitatory/inhibitory (E/I) balance [33]. Pyr and PV neurons have different intrinsic biophysical properties: in terms of excitability, PV cells have a higher threshold and gain compared to Pyr neurons [34]. These properties, along with other neuronal characteristics, such as refractory period and membrane time constants, contribute to the characteristic regular-spiking patterns of pyramidal cells and fast-spiking patterns of PV interneurons.

In our hardware experiments, we reproduce the properties of both Pyr and PV neuron types on a mixed-signal analog/digital Dynamic Asynchronous Neuromorphic Processor, called DYNAP-SE2 [31] (see Fig. 1a), which implements a silicon neuron circuit equivalent to the adaptive exponential integrate-and-fire neuronal model. The dynamics of the different neuron classes were reproduced by tuning the refractory period, time constants, spiking threshold, and neuron gain parameters (Table 5, in the Supplementary Material) to match the corresponding biologically measured values (Fig. 1b). Since parameters are shared by all neurons in a chip core, we assign each neuron population type to a different core (Fig. 1a). Even though all neurons in a core share the same parameters, variations in their intrinsic characteristics arise as a result of analog chip mismatch. For instance, one can observe differences between two excitatory or inhibitory neurons recorded within the same core, as illustrated in Fig. 1c. Overall, the neural dynamics and Input-Frequency (FI) curve of recorded excitatory and inhibitory neurons on chip (Fig. 1c-d) show that the choice of parameters used for the silicon neurons is compatible with the behavior of biological PV and Pyr cells [34]. In the following section, we employ cross-homeostatic plasticity [28] to self-tune the connections between PV and Pyr populations so that spontaneous activity in the inhibition-stabilized regime emerges in the network.

**Figure 1:**
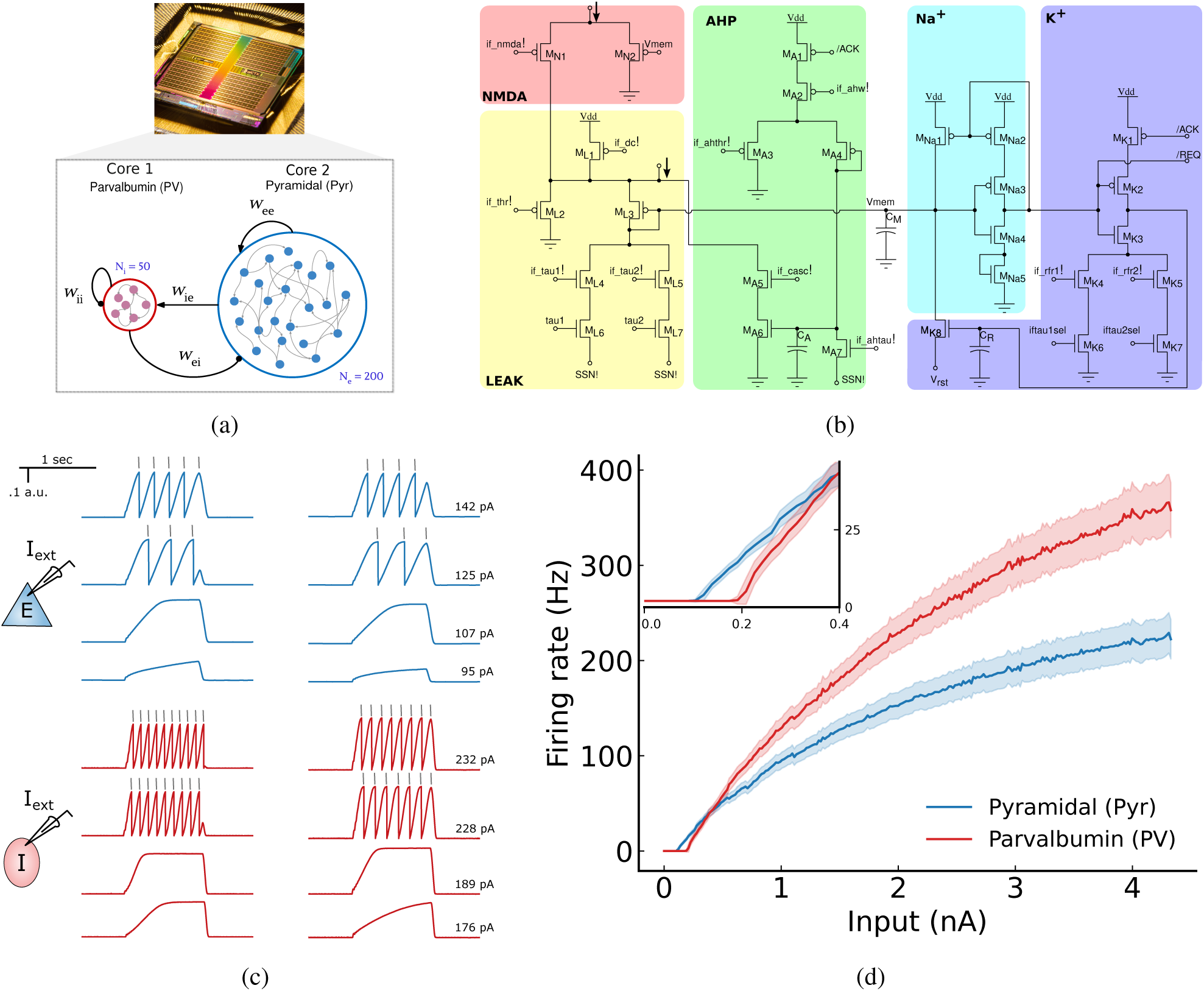
(a) Image of the DYNAP-SE2 chip, showing the four cores. An E-I network is implemented on chip. Excitatory (Pyr) and Inhibitory (PV) neurons occupy different cores, and are connected by four weight classes, Pyr-to-Pyr (𝑤_𝑒𝑒_), Pyr-to-PV (𝑤_𝑖𝑒_), PV-to-PV (𝑤_𝑖𝑖_), and PV-to-Pyr (𝑤_𝑒𝑖_). (b) Circuit diagram depicting a DPI neuron, with each component within the circuit replicating various sub-threshold characteristics (such as adaptation, leakage, refractory period, etc.) inherent to an adaptive exponential neuron [31]. (c) Neuron membrane potential traces over time as a response to DC injection. Blue traces show two example cells from the excitatory chip core in response to four current levels. Red traces show two sample PV cells from the inhibitory chip core. The relationship between the measured voltage and the internal current-mode circuit output that represents the membrane potential is given by an exponential function that governs the subthreshold transistor operation mode [32]. In each case, the bottom two traces show neuronal dynamics in the subthreshold range. The top two traces show neuronal dynamics as the membrane potential surpasses the spike threshold (spike added as grey line to indicate when the voltage surpasses threshold). DC injection values are indicated on every trace. (d) Input-response curves for the resulting Pyr and PV neurons on-chip. Inset: zoomed-in view at low firing rates. In (c), it is evident that a higher input current was required to elicit spikes in PV neurons. This phenomenon can be attributed to the higher threshold of PV neurons.

### The network converges to stable self-sustained dynamics

As an overview of the experimental paradigm, the training procedure begins with random initialization of the network weights. Then, at each iteration, we record the resulting firing rates, calculate the weight update following the cross-homeostatic equations for the four weight classes (𝑤_𝑒𝑒_, 𝑤_𝑒𝑖_, 𝑤_𝑖𝑒_, and 𝑤_𝑖𝑖_, equation 3, see Methods) and apply the new weight values to the chip.

Fig. 2a illustrates an example of the evolution of firing activity during the course of training for Pyr and PV cells. Starting from a random value determined by the initialization of weights, the average rates converge to their target values. Fig. 2b plots the evolution of weights during training. On chip, the weights are controlled by two parameters, a coarse 𝐶_𝑤_ and a fine 𝐹_𝑤_ value (See Methods, eq. 5). On most iterations, only the fine value 𝐹_𝑤_is updated, but updates to the coarse value 𝐶_𝑤_ occur occasionally. Even though the coarse updates cause abrupt jumps in the weight value (due to the nature of the bias-generator circuit implementation), the cross-homeostatic rule is robust enough to guide the network to a stable regime. Fig. 2c shows a raster plot from the network in Fig. 2a-b at a converged state. For an additional example with different weight dynamics see Fig. S 10. It should be emphasized that, while an initial “kick” was provided to the network to engage recurrent activity (40 ms at 100 Hz), no further external input was introduced during the trial; hence, any observed spiking activity originates entirely from internal mechanisms.

**Figure 2:**
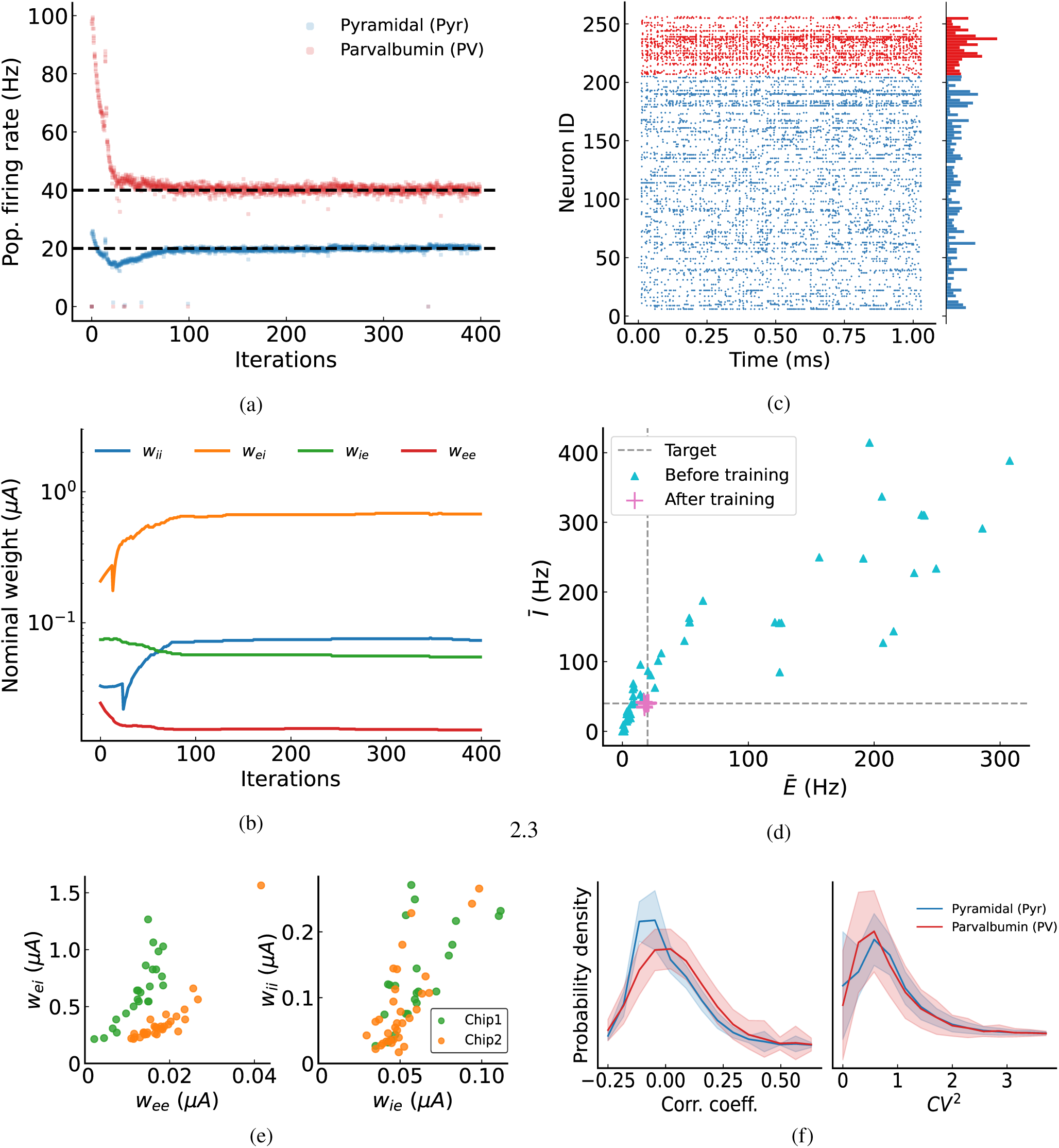
(a) A representative experiment of the convergence of excitatory (red) and inhibitory (blue) firing rates during learning. The gray dashed lines show the desired set-point targets. (b) A representative experiment (the same as in panel a) of the convergence of weight values during learning. The plotted value is nominal, inferred from eq. (1) (see Methods). (c) Raster plot exemplifying on-chip firing activity during a single emulation run at the end of training. Both Pyr and PV neurons successfully converge to an asynchronous-irregular firing pattern at the desired population FRs. (d) Initial and final firing rates across different chips (n=57, across two chips) and different initial conditions. For each initial condition, network connectivity was randomized. (e) Left: relationship between the final 𝑤_𝑒𝑒_ and 𝑤_𝑒𝑖_ values across different iterations. Orange and green colors correspond to two different chips. Right: same for 𝑤_𝑖𝑖_ and 𝑤_𝑖𝑒_. (f) Distributions for the 5 ms correlation coefficients between neurons (left) and coefficient of variation (𝐶𝑉 ^2^) (right), demonstrating low regularity and low synchrony. The line indicates the mean. Shaded region indicate standard deviation.

When repeating the training process from different initial conditions of randomly sampled weights (see Fig. S 9 for the corresponding initialization), the weights converge to different values, which however produce the same desired network average firing rate [28]. Fig. 2d illustrates the rate space with multiple initialization and convergence to respective target set-points (dashed lines). The root mean square error from the target is 0.945 Hz for 𝐸and 1.653 Hz for 𝐼. The converged weight values are approximately aligned to a linear manifold (Fig. 2e),

where the sets of excitatory and inhibitory weights 𝑤_𝑒𝑒_, 𝑤_𝑒𝑖_ and 𝑤_𝑖𝑒_, 𝑤_𝑖𝑖_, are correlated (correlation coefficients and p-values are as follows: 𝑤_𝑒𝑒_ and 𝑤_𝑒𝑖_: 𝜌_𝑔𝑟𝑒𝑒𝑛_ = 0.81 and 𝑝 = 1.55 · 10^−06^; 𝜌_𝑜𝑟𝑎𝑛𝑔𝑒_ = 0.88, 𝑝 = 6.6 · 10^−12^; 𝑤_𝑖𝑒_ and 𝑤_𝑖𝑖_: 𝜌_𝑔𝑟𝑒𝑒𝑛_ = 0.63 𝑝 = 0.0009; 𝜌_𝑜𝑟𝑎𝑛𝑔𝑒_ = 0.75, 𝑝 = 3.9 · 10^−07^, where *green* and *orange* refer to two different chips). This is well aligned with the theoretical solution derived for rate E-I networks at their set-points, when the neuron transfer function is linear or threshold-linear [28, eqn. 4-5]. In our case, the linear transfer function assumption does not hold, because the AdEx silicon neuron models saturate at a rate that is inversely proportional to the refractory period parameter. However, the linear approximation holds well in the region of operation forced by the learning rule, set at relatively low firing rates (see Fig. 1d).

### The network is in an inhibition-stabilized asynchronous-irregular firing regime

Next we investigated the properties of the final network dynamics after learning. Fig. 2c shows a sample of activity of the converged network, as recorded from the chip, in which the excitatory and inhibitory populations exhibit asynchronous-irregular activity patterns. The minimal correlations among neurons (averages ⟨𝜌 _𝑝𝑦𝑟_ ⟩ = 0.037, ⟨𝜌 _𝑝𝑣_⟩ = 0.069) serve as evidence of their asynchronous activity (Fig. 2f, left). Irregularity is shown by the coefficient of variation (CV^2^) of most neurons being close to 1 (averages ⟨𝐶𝑉 ^2^ ⟩ = 0.949, ⟨𝐶𝑉 ^2^ ⟩ = 0.945) (Fig. 2f, right) . The CV^2^is computed as the variance of the inter-spike interval divided by its squared mean, and equals 0 for perfectly regular firing, and 1 for Poisson firing [35]. It is worth noting that during the 1-second simulations, we do find instances of synchronization, which can be attributed to the small population size of the network and increased firing rate in comparison to the baseline spontaneous activity. As an average however, we conclude that cross-homeostatic plasticity brings the network to an asynchronous-irregular firing regime, what is typical in realistic cortical networks [36].

As further evidence of the inhibition-stabilized regime of the network, we demonstrate the occurrence of the “paradoxical effect” in our deployed network on chip [18, 20]. The paradoxical effect is a hallmark of inhibition-stabilized networks that has been observed in cortical circuits. When inhibitory neurons are excited (either optogenetically *in vivo*, or via an external current in computational models) their firing rates show a paradoxical decrease in activity during the stimulation, providing clear evidence that that the network is in an inhibition-stabilized regime (Fig. 3a). Analogously, when we inject an external depolarizing pulse to all PV neurons on chip for 0.2 s we observe a decrease in the firing rate of both cell populations for the duration of the stimulus, returning back to the target FR when the stimulus ends (Fig. 3b). Across all 14 iterations we use pairwise t-tests to assess the impact of stimulating PV neurons. For PV neurons, stimulation significantly reduces activity during stimulation compared to pre- and post-stimulation states (𝑝 < 10^−18^ in both cases). Conversely, there is no significant difference between pre- and post-stimulation activity, indicating recovery. Five trials, which were excluded from the analysis, do not exhibit recovery because the activity is entirely suppressed by the strength of the paradoxical effect. A similar trend is observed for Pyramidal neurons (𝑝 < 10^−20^).

**Figure 3:**
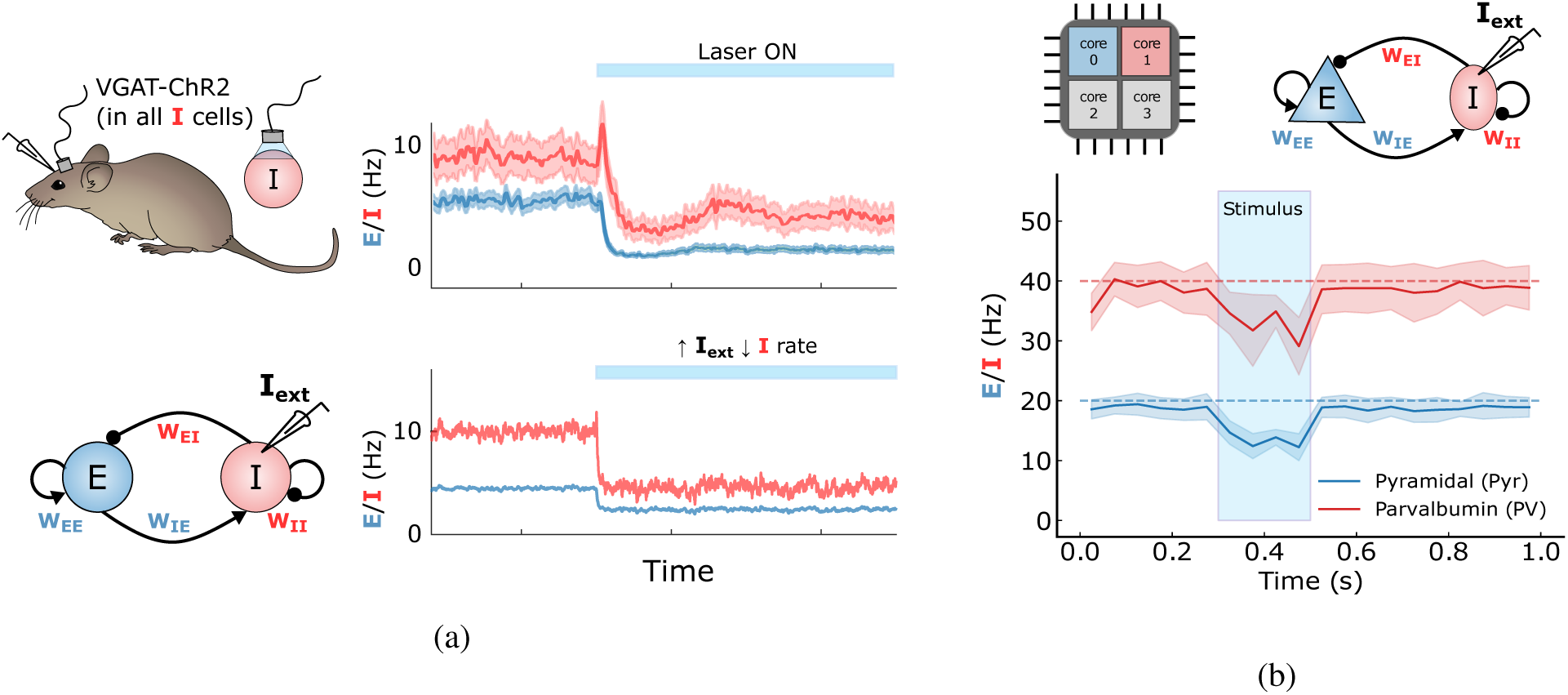
(a) The paradoxical effect is a well known phenomena in cortical circuits. In the awake resting cortex of mice (upper figure), when inhibitory neurons are optogenetically activated, their firing rates (in red) show a paradoxical decrease in activity during the stimulation, indicative of an inhibition-stabilized network. Adapted from Sanzeni et al. [21]. Similar to the experimental case (bottom figure), numerical simulations of firing rate models with sufficient excitatory gain and balanced by inhibition show the paradoxical effect when the inhibitory population is excited via an external current. (b) The paradoxical effect can be demonstrated in analog neuromorphic circuits by applying a Poisson input of 250 Hz for 200 ms to the inhibitory units of networks converged to self-sustained activity via cross-homeostasis. The resulting decrease in the firing rate of the inhibitory units demonstrates that the on-chip network is in the inhibition-stabilized regime. No. of trials = 14, shading region represents standard deviation in firing rate across trials.

### Inhibitory plasticity is necessary to achieve reliable convergence

The equations for the weights in absence of noise [28, eqn. 4-5] and the experimental results in Fig. 2 e indicate that two of the four weight parameters, for example, the inhibitory weights 𝑤_𝑖𝑖_ and 𝑤_𝑒𝑖_, can be chosen arbitrarily, since one can always find a solution by learning the other two. This appears to suggest that homeostatic plasticity of the inhibitory weights might not be required in an ideal case. However, we experimentally show that excitatory plasticity alone is not sufficient to reliably make the network converge in the presence of the noise and non-idealities of the analog substrate (see Fig. 4). In the absence of inhibitory plasticity, both the excitatory and inhibitory populations approach their respective set-points, but they fail to converge to stable activity levels. In addition, the network is in an unstable configuration, as can be seen by the many “exploding” trials that occur during the experiment, where the firing rate diverges and is limited only by the saturation of the spiking dynamics. (Fig. 4a). In contrast, the full cross-homeostatic plasticity rule robustly drives the network activity towards the set-points after just a few iterations (Fig. 2a). These results hold for multiple initializations (Fig. 4b), indicating that inhibitory plasticity is necessary for robust convergence.

**Figure 4:**
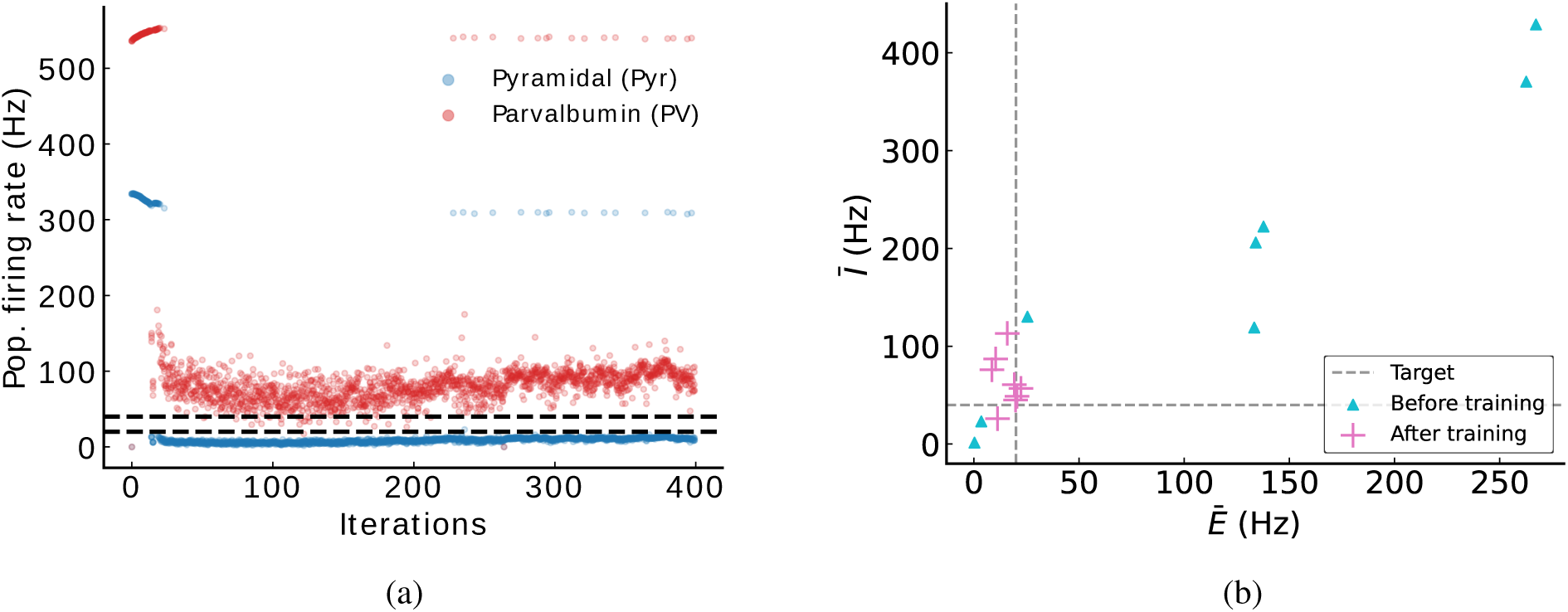
(a) Example network with training results on “fixed” inhibitory synapses. Convergence is impaired if inhibitory plasticity is turned off. Note that many trials at the end of the experiment exhibit “exploding” rates above 300 Hz, corresponding to the neuron’s saturation rates. (b) Random networks ran with different initialization of all weights (n=8). The firing rates do not converge to the targets as precisely as in Fig. 2a, having a much higher root mean square error from the target (RMS on 𝐸: 6.39, on 𝐼: 35.11).

Homeostatic plasticity in inhibitory neurons is less studied than in excitatory neurons [24, 37], although there is evidence that inhibitory neurons also regulate their activity levels based on activity set-points [38, 39]. Considering our on-chip results, we therefore speculate that homeostatic plasticity in all excitatory and inhibitory connections plays an important stabilization role also in biological systems.

### Excitatory plasticity is desirable for reliable convergence and chip resource utilization

Given that inhibitory plasticity seems to be crucial, we next investigated whether excitatory-to-excitatory plasticity is also necessary, providing the fixed value of 𝑤_𝑒𝑒_ is sufficiently high to generate self-sustained activity. Having𝑤_𝑒𝑒_ free from any homeostasis could be beneficial when introducing other forms of plasticity, such as standard associative learning rules (eg. Hebbian learning or spike timing dependent plasticity (STDP)). We thus ran a set of experiments with 𝑤_𝑒𝑒_ plasticity frozen. We find that in most cases, the network is able to converge to stable activity levels at the setpoints (Fig. S 11). However, because of the fewer degrees of freedom, the rules fail to converge whenever any of the weights reaches their maximum possible value on chip (Fig. S 11b). We conclude that although it is possible to operate cross-homeostasis without 𝑤_𝑒𝑒_ plasticity, it is highly beneficial to maintain all four weight classes plastic for robust convergence across all initial conditions. This is true also when freezing weights other than 𝑤_𝑒𝑒_. To confirm these results, we ran multiple simulations on a rate model (see Fig. S 12), freezing different sets of weights. Indeed, our results show that having plasticity in all weight classes allows for robust convergence across a wide range of initial conditions, while such convergence is always partly impaired whenever plasticity is blocked on any weight class.

Despite the above evidence that excitatory-to-excitatory plasticity is highly beneficial, these results do not rule out that alternative forms of homeostasis could still succeed in bringing the network to fully self-sustained regimes without operating on 𝑤_𝑒𝑒_. Recently, a set of homeostatic rules operating at the 𝑤_𝑒𝑖_ and 𝑤_𝑖𝑒_ weights has been proposed which implement stimulus-specific feedback inhibition consistent with V1 microcircuitry [40]. The rules proposed by Mackwood and colleagues are simple enough that they can be implemented with our chip-in-the-loop setup, as are the cross-homeostatic plasticity rules (see Methods). In fact, the rule for the 𝑤_𝑖𝑒_weight is the same as its cross-homeostatic counterpart. The rule for 𝑤_𝑒𝑖_ obeys a more classic homeostatic formulation [24, 41]. Unlike cross-homeostasis, these rules only impose an activity setpoint for the excitatory population. We decided to test whether, in the absence of external input but with sufficiently high 𝑤_𝑒𝑒_, the rules from Mackwood and colleagues could be employed to generate self-sustained dynamics in the inhibition-stabilized regime on chip. The experiments show that, in most cases, the network converges with stable self-sustained dynamics to the established excitatory setpoint (Fig. 5 a- b). However — as happened when dropping plasticity in some of the cross-homeostatic weights — because of the lower degrees of freedom, the rules fail to converge whenever any of the weights reaches their maximum possible value on chip (Fig. S 11b). Another disadvantage of employing these rules is the lack of inhibitory setpoint for the I population. Although this could be perceived as advantageous (to keep inhibitory rates free of constraints), in practice the network often converges to very large values of inhibitory rates (in most cases above 100 Hz) (Fig. 5 c). These high rates are often un-biological, and not desirable for neuromorphic applications where sparsity and ultra-low power consumption are a constraint.

**Figure 5:**
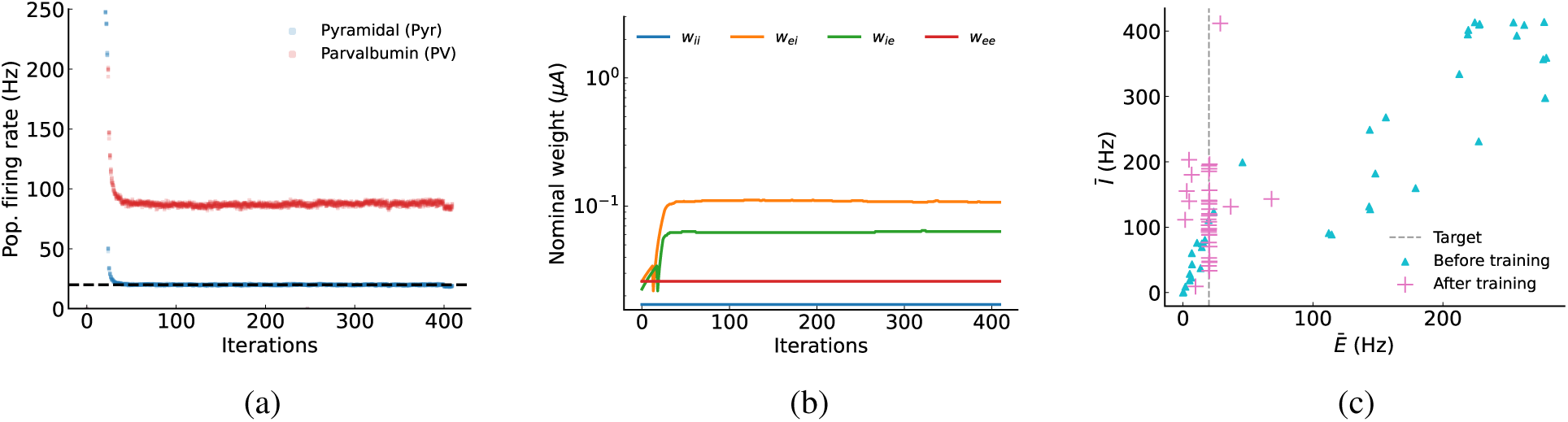
Results obtained using Mackwood et al.’s rule. (a) convergence of the firing rate to stable self-sustained dynamics. Note that, in this case, there is no inhibitory target. (b) convergence of the weight values. 𝑤_𝑖𝑖_ and 𝑤_𝑒𝑒_are fixed and thus do not show any dynamics under this rule. (c) initial and final firing rate values over many experimental runs. 𝐸 mostly converges to the target, while, unlike with cross-homeostasis, the final 𝐼 is left to vary freely (n=34).

In light of these results we envision cross-homeostatic plasticity as a powerful tool to robustly calibrate neuromorphic hardware, likely after fabrication, into a fully self-sustained inhibition stabilized regime. After initial calibration — which would bring the four weights to a stable low-firing regime across a broad range of initial conditions — the experimenter could choose, for example, to freeze homeostasis on the excitatory weights, or use the rules proposed by Mackwood and colleagues, to allow for more freedom to other plasticity forms.

### Multiple subnetworks can be emulated on a single chip

In the network discussed so far, we modeled a single stable fixed point attractor in firing rate space, using an E/I-balanced network. We used 200 out of the 256 neurons available on a single core of our multi-core neuromorphic chip (DYNAP-SE2).

Even though it is still a relatively small-scale network, for computational purposes, having a single attractor formed from 200 neurons is not an optimal utilization of the resources on the chip. Employing smaller networks would allow us to implement multiple attractors on a single chip, whereby each subnetwork could represent, for example, a distinct choice in a decision-making task. This could be useful for stateful computations, or for example, to implement different recurrent layers on a chip [42].

Therefore, we also run scaled down variations of the previous network, and assess whether cross-homeostatic plasticity can still bring activity to desired levels. Table 1 summarizes the population count, connectivity scaling, and adjustment in the target firing rate that we adopt for each experiment.

**Table 1:** Variation in network size, connection scaling, and stable target firing rate (set-points) for reliable self-sustained activity.

| Number of neurons (E/I) | Connection Probability | Target (E/I) |
| --- | --- | --- |
| 200/50 | 0.1 | 20/40 Hz |
| 100/25 | 0.1 | 20/40 Hz |
| 64/16 | 0.35 | 25/50 Hz |
| 48/12 | 0.35 | 40/60 Hz |

Furthermore, we experiment with implementing multiple copies of these scaled-down networks on the same chip. We create 5 instances of different E/I networks, each with 50 excitatory and 15 inhibitory neurons. The connectivities within each sub-network are drawn from the same distribution. Each subnetwork does not cross-talk with others. The configuration is illustrated in Fig. 6a. To implement this configuration on the chip, we group the excitatory neuron populations into a single core, and use another core for the inhibitory populations. All parameter values, including weights, are shared between sub-networks. The challenge is to check if homeostatic plasticity leads to convergence for all subnetworks, since the effect of the weights is different for each E/I network due to the inhomogeneity of the chip.

**Figure 6:**
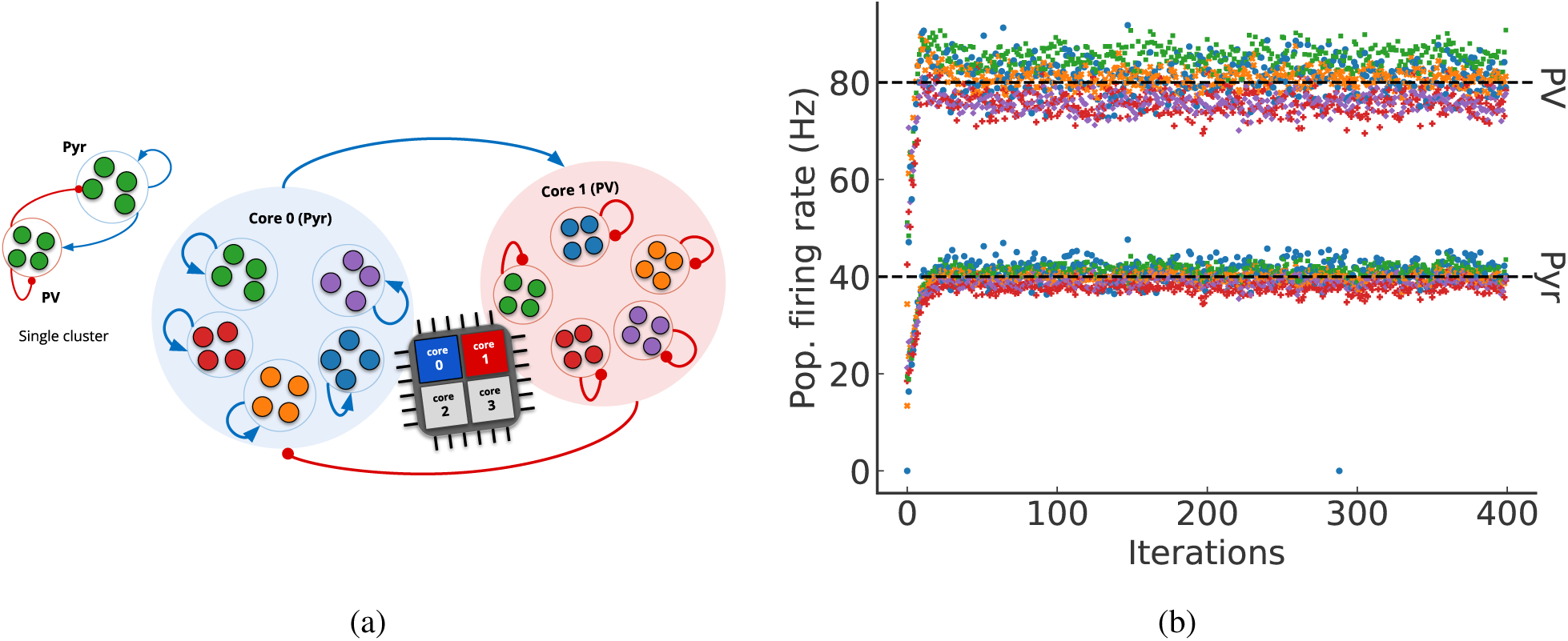
Simultaneous training of multiple clusters. (a) Schematic of clusters on the chip. One core contains excitatory populations, and another contains inhibitory populations. We implemented five E/I networks, with each network comprising 50 excitatory and 15 inhibitory neurons. As per table 1, the connection probability within these networks stands at 0.35. To account for the network’s scaling, we defined higher set points. It’s worth noting that these clusters do not interconnect with one another. b) Rate convergence for 5 clusters sharing the same nominal weights. Each color represents one cluster.

During the training process, we sequentially stimulate one sub-network at a time during each iteration. We then compute the change in synaptic weights, denoted as Δ𝑤, based on the activity of the neuron population within that E/I network. The weight is configured on the chip after each iteration based on the average Δ𝑤 across all sub-networks. Fig. 6b shows that all five subnetworks, with shared weights, can indeed converge to mean firing rate regimes close to the target firing rate.

### Neuronal ensembles remain intact under cross-homeostasis

The effect of homeostasis is to maintain a set level of activity in neurons by scaling synaptic efficacy, as done in this work, or by changing the intrinsic properties of the neuron. This raises concerns about the stability-plasticity dilemma, i.e., the competition between the homeostatic stability mechanism and the associative forms of synaptic plasticity that contribute to the formation of functional *neural ensembles*, as they may counteract each other causing either forgetting or the inability to form new memories. A neural ensemble corresponds to a group of interconnected neurons (e.g., that increased their synaptic weights via learning) that contributes to the representation of a particular feature or concept. Typically, neurons forming an ensemble exhibit shared selectivity [43, 44].

The ensemble could potentially be affected by the homeostatic scaling process, which attempts to equalize the activity levels across the network. To verify that the cross-homeostatic plasticity used here does not interfere or disrupt the role of neural ensembles we artificially formed a “crafted memory” into the network during the homeostatic training process (Fig. 7). This memory is implemented as a neural ensemble within the excitatory population of the network. Specifically, the ensemble consists of 32 neurons, in which each neuron has a connection probability with other neurons of the same ensemble of 0.5, five times more than the rest of the network (Fig. 7a).

**Figure 7:**
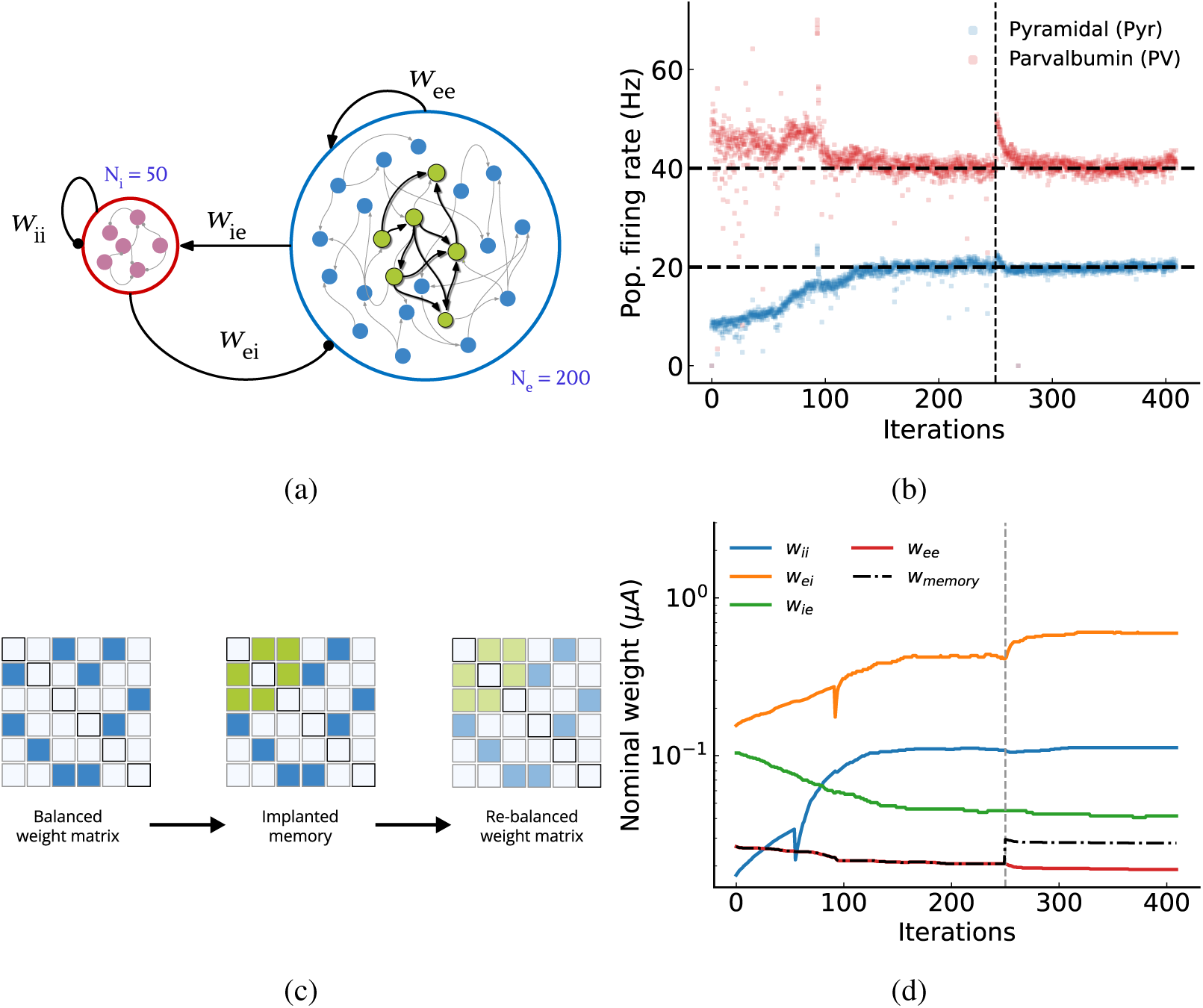
Training result with a memory implanted in the network. (a) Random recurrent E/I network with a neural ensemble (size = 32, green neurons). (b) Convergence of both excitatory (Pyr) and inhibitory (PV) populations to their respective set-point. A subset of the weights is manually changed to represent an ensemble at iteration 250, well after the network has converged to its set-point. A jump in activity is observed in both Pyr and PV cells due to implanted memory. (c) Diagram illustrating the process of rebalancing the weight matrix after implanting a single memory. Due to the chip’s limited observable and shared parameters, it’s not possible to record the actual weight matrix for visualization. (d) The weight (𝑤_𝑒𝑒_) value for 32 neurons is increased by a factor of 0.0088 µA, illustrated by a dotted line. All weights undergo alteration again to compensate for the ensemble, bringing the network back to the set point after the memory is implanted. However, the memory remains intact, as shown by higher recurrent weight (𝑤_𝑚𝑒𝑚𝑜𝑟 𝑦_) compared to the baseline excitatory connectivity (𝑤_𝑒𝑒_).

After 250 cross-homeostatic plasticity training iterations, we manually form the ensemble by introducing the additional connections in it. We observe an immediate increase in the overall activity of both excitatory and inhibitory neurons (Fig. 7b) after ensemble implantation, which is however reduced as the homeostatic training continues. The change in weights (Fig. 7d-c) illustrates how inhibitory plasticity (specially 𝑤_𝑒𝑖_) compensates for increased activity, bringing the firing rates back to the target point. Because the values of weights within and outside of the ensemble remain differentiated, even after homeostatic plasticity converges, the neural ensemble remains intact and retains its selectivity for the specific feature it represents. We conclude that the cross-homeostatic learning rule brings the activity back to the target while preserving the implanted memory.

### Multiple stable ensembles can coexist with emergent sWTA dynamics

The previous results show that a single memory could be implanted long-term in the presence of cross-homeostasis. We extended the model as a proof-of-concept to craft multiple memory clusters with inter-cluster connectivity, and demonstrate to what extent they could be individually recalled. Fig. 8 a- c illustrates a network with three stable implanted memories, as described in the previous section. These individual memories can be robustly recalled even when inputs are presented to other memory clusters. Remarkably, these implanted memories exhibit emergent soft-winner-take-all (sWTA) dynamics, where the neural population receiving the strongest stimulus “wins” and exhibits an increased firing rate during recall. This cross-cluster competition arises from lateral inhibition between clusters, as demonstrated in rate-based network simulations. Surprisingly, this occurs despite no specific inhibitory motifs being crafted into the network, unlike in previous WTA models. [12–15].

**Figure 8:**
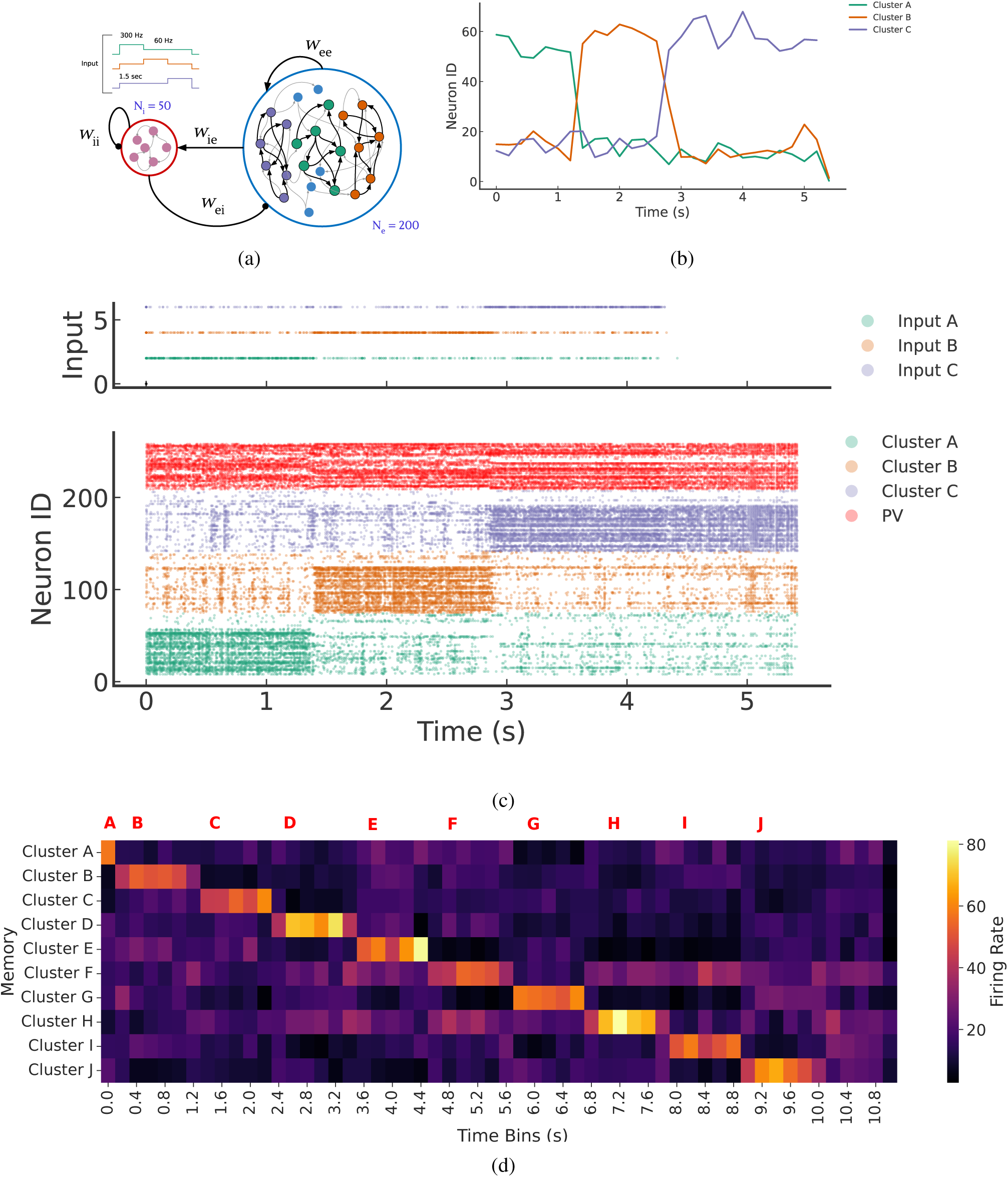
Winner-take-all dynamics with three sub-clusters implanted in the network. (a) Random recurrent E/I network with three neural ensembles (size = 50, green, orange and purple neurons). The simulation protocol (top left) shows how different sub-clusters receives inputs over the course of simulation. (b) Firing rates of all clusters when each cluster is stimulated with either high (300 Hz) or low frequency (60 Hz) inputs. The cluster receiving the high-frequency input dominates the competition. We demonstrate that no special tuning is necessary to achieve this winner-takes-all (WTA) phenomenon. (c) Raster plot illustrating spike activity for all neurons. For the final stimulation, the plot shows that the cluster remains active even after the stimulus is withdrawn, showing a form of working memory. (d) Extended version of the memory network with 10 sub-clusters, each with 20 neurons.

Cross-homeostasis therefore implicitly implements sWTA dynamics into recurrent networks. Interestingly, we observe that some memory clusters also exhibit working memory properties, as the increase in firing rate upon input activation is maintained once the input is withdrawn from the network Fig. 8 b- c. While such working memory properties are not present in all clusters, we have also observed them in simulations of rate-based networks (see Fig. S 13). Further research should explore the conditions under which persistent activity is maintained after input withdrawal and examine the robustness of this activity in the face of external perturbations.

We finally show that the chip supports the implementation of up to 10 coexisting memory clusters that can be robustly recalled independently (Fig. 8 d). These networks could be used to implement decision-making processes on chip, along with other interesting computations such as signal-restoration or state-dependent processing. We conclude that cross-homeostatic plasticity not only allows for maintenance of information, but imposes, “for free”, computationally relevant dynamics into the network.

## Discussion

Hardware implementations of spiking neural networks are being proposed as a promising “neuromorphic” technology that can complement standard computation for sensory-processing applications that require limited resource usage (such as power, memory, and size), low latency, and that cannot resort to sending the recorded data to off-line computing centers (i.e., “the cloud”) [30, 45, 46]. Among the proposed solutions, those that exploit the analog properties of their computing substrate have the highest potential of minimizing power consumption and carrying out “always-on” processing [9, 47]. However, the challenge of using this technology is to understand how to carry out reliable processing in the presence of the high variability, heterogeneity and sensitivity to noise of the analog substrate. This challenge becomes even more daunting when taking into account the variability of memristive devices, which are expected to allow for the storage and processing of information directly on chip [48].

In this paper, we show how resorting to strategies used by animal brains to solve the same challenge can lead to robust solutions: we demonstrated that cross-homeostatic plasticity can autonomously bring analog neuromorphic hardware to a fully self-sustained inhibition-stabilized regime, the regime that is thought to be the default state of the awake neocortex [19–22]. These are the first results to show how biologically inspired learning rules can be used to self-tune neuromorphic hardware into computationally powerful states while simultaneously solving the problem of heterogeneity and variability in the hardware.

Our results also validate the robustness of the cross-homeostatic plasticity hypothesis [28] in a physical substrate that presents similar challenges to those of faced by biological networks. We also highlight the crucial contribution of inhibitory plasticity, specifically, that homeostatic plasticity of inhibitory connections is necessary to maintain stability in both biological and silicon networks. We further show that although not necessary, excitatory homeostatic plasticity is desirable to guarantee convergence across a wide range of initial conditions. Additionally, by comparing our results with those given by using a different learning rule [40], we demonstrate that having a setpoint for the inhibitory population is also beneficial, in order to keep firing rates into a biological regime. These low firing rates are also technologically relevant, as neuromorphic applications benefit from the sparsity and ultra-low power consumption of networks in this regime.

Notably, we constructed an on-chip stable attractor neural network that effectively maintains a long-term memory represented by a neural ensemble, and demonstrates that multiple networks, each exhibiting fixed-point attractor dynamics, can coexist within the hardware [18, 36]. We next show that this stable multi-network implementation presents emergent sWTA dynamics, even though no specific connectivity motives have been introduced in the network, unlike in previous work [12–15]. Networks with sWTA dynamics can perform powerful computations, ranging from signal-restoration to state-dependent processing. Cross-homeostatic plasticity could thus be used to configure networks with self-generated dynamics that are thought to underlie a number of computations including decision-making, motor control, timing, and sensory integration [49–51].

By taking inspiration from the self-calibration features of biological networks, we achieve a remarkable level of resilience to hardware variability, enabling recurrent networks to maintain stability and self-sustained activity across different analog neuromorphic chips. Cross-homeostatic plasticity allows us to instantiate neural networks on different chips without the need for chip-specific tuning, thereby addressing the inherent mismatch in analog devices. Such bio-plausible algorithms provide a promising avenue for the development of robust and scalable hardware implementations of recurrent neural networks, while mitigating the challenges associated with the intrinsic mismatch in analog devices.

In short, these results not only provide insights into the underlying mechanisms configuring biological circuits, but also hold great promise for the development of neuromorphic applications that unlock the power of recurrent computations. This research presents a substantial step forward in our understanding of neural dynamics and paves new possibilities for practical deployment of neuromorphic devices that compute, like biological circuits, with extremely low-power and low-latency using attractor dynamics.

## Methods

### Neuromorphic hardware: the spiking neural network chip

In this study, we use a recently-developed mixed-signal spiking neural network chip, called Dynamic Neuromorphic Asynchronous Processor (DYNAP-SE2) [52]. This chip comprises 1024 adaptive-exponential integrate-and-fire (AdEx I&F) neurons [8, 53], distributed across four cores. The neuron parameters consist of its spiking threshold, refractory period, membrane time constant, and gain. On hardware, the inputs to each neuron are limited to 64 incoming connections, implemented using differential pair integrator (DPI) synapse circuits [54]. Each synaptic input can be configured to express one of four possible temporal dynamics (via four different DPI circuits): AMPA, NMDA, GABA-A, and GABA-B. Besides expressing different time constants, the excitatory synapse circuits (AMPA and NMDA) differ in that the NMDA circuit also incorporates a voltage gating mechanism. Similarly, the inhibitory synapses (GABA-A, and GABA-B) differ in their subtractive (GABA-B) versus shunting (GABA-A) effect.

The bias parameters that determine the weights, neuronal and synaptic properties are set by an on-chip digital to analog (DAC) bias-generator circuit [55] and are shared globally within each core. These parameters are expressed as currents that drive the neuromorphic circuits and are represented internally through a “coarse” and a “fine” value, according to the following equation:

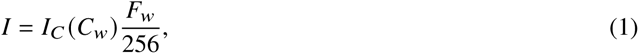

where 𝐹_𝑤_ represents the fine value and 𝐶_𝑤_ the coarse one. The variable 𝐼_𝐶_ represents a current estimated from the bias-generator circuit simulations, which depends on 𝐶_𝑤_ as specified in Table 2.

**Table 2:** Nominal estimated coarse current values of the on-chip bias generator circuit for synaptic weights.

| $C_w$ | 0 | 1 | 2 | 3 | 4 | 5 |
| --- | --- | --- | --- | --- | --- | --- |
| $I_C$ (nA) | 0.07 | 0.55 | 4.45 | 35.0 | 280 | 2250 |

Before training, the network is initialized with 𝐶_𝑤_ randomly taking a value of 3, 4, or 5, and 𝐹_𝑤_ uniformly distributed between 20 and 200. The corresponding currents will therefore range between 2.7 nA and 1757 nA. We avoided using lower values of 𝐶_𝑤_ because the neuron remains inactive when stimulated with such small current injections. Additionally, initializing all weights with very high values causes the chip to produce too many spikes for the digital interface to the data-logging computer to handle.

Due to device mismatch and variability in the analog circuits, even when many circuits share the same nominal biases, the effective values will vary. On the DYNAP-SE2 and other mixed-signal neuromorphic processors implemented using similar technology nodes and circuit design styles, the coefficient of variation of these parameters is typically around 0.2 [10]. The learning rules proposed in this work make these networks robust to variability with these figures, and lead to a stable activity regime.

### Network emulation

We create an on-chip E/I network composed of 200 Pyr (excitatory) and 50 PV (inhibitory) neurons. Each neuron type is implemented by tuning the refractory period, time constants, spiking threshold, and neuron gain parameters to match biologically measured values (Fig. 1c, [34]). The two populations are sparsely connected with a 10% connection probability for each pair of neurons (both, within population and between populations). Synaptic weights take four sets of values shared globally among all Pyr-to-Pyr (𝑤_𝑒𝑒_), Pyr-to-PV (𝑤_𝑖𝑒_), PV-to-PV (𝑤_𝑖𝑖_), and PV-to-Pyr (𝑤_𝑒𝑖_) connections respectively. All excitatory weights represent synapses modeled as AMPA receptors, and inhibitory weights as GABA-A (somatic). Here, 𝑤_𝑖𝑖_ and 𝑤_𝑒𝑖_ are somatic inhibitory connections. Even though the weight parameters are shared, the synaptic efficacy of each synapse is subject to variability due to device mismatch in the analog circuits [10].

During training, synaptic weights are adjusted such that the average activity for both Pyr and PV population converge to their respective target firing rates (or *set-points*), and can thus maintain self-sustained activity in the absence of any external input or bias current. To kickstart activity in the network, we provide a brief 40 ms external input of four spikes at 10 ms intervals to 80% of the Pyr cells, with random delays for each postsynaptic target neuron to introduce temporal sparsity to the initial response of the population. The network is emulated in real-time on the chip (i.e., the dynamics of the analog circuits evolve through physical time reproducing the dynamics of the biological neural circuits). Each experiment is run for 1 s and the spiking activity of the whole network is streamed to a computer, for data logging and analysis purposes.

In the analysis, the network response is measured by calculating the average firing rate of both neural populations, which is then used to calculate the error (i.e., the difference between set-point and actual activity) that drives the weight update (see next section). As we are interested in the network’s self-sustained behavior, we discard the response induced by the transient external input while calculating the firing rates (i.e., we ignore the first 60 ms of activity, which comprises the first 40 ms of external stimulation and an additional stabilization window of 20 ms). During the initial stages of the training procedure, before E/I balance is reached, the network could generate non-sustained bursts of activity. Therefore, we calculate the firing rate by considering only the time window when the neural populations are active, i.e. the in-burst firing rate, as opposed to the average over the whole trial. We empirically find that this choice leads to the desired sustained activity.

### Learning rule

To bring the recurrent dynamics to a fully self-sustained inhibition-stabilized regime we employ the recently proposed cross-homeostatic plasticity rule [28], which tunes all four synaptic weight classes in parallel.

In classic forms of homeostatic plasticity, both excitatory and inhibitory populations self-regulate calcium levels based on their own activity [24, 37]. In contrast to these classic forms, where a neuron tunes its weights to maintain its own average output of activity (a firing rate set-point), the cross-homeostatic weight update rule aims at bringing the neuron’s presynaptic partners of opposite polarity to an average population firing rate setpoint, according to the following equations:

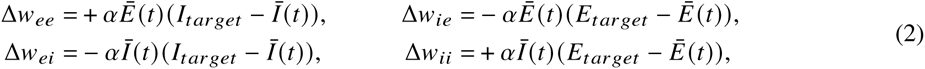

where 𝛼 is the learning rate (𝛼 = 0.05), 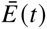 and 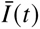 are the measured population firing rates averaged over 1s during a trial 𝑡 and 𝐸_𝑡𝑎𝑟𝑔𝑒𝑡_ = 20 Hz, 𝐼_𝑡𝑎𝑟𝑔𝑒𝑡_ = 40 Hz are the desired firing rate set-points, for the Pyr and PV populations respectively. This type of plasticity can be interpreted biologically as having a target on the input current that a given neuron receives. For example, an excitatory neuron could have indirect access to the average firing rate of their presynaptic inhibitory neurons based on a running average of the total GABA it receives.

While biologically plausible target activities are typically lower (e.g., 5 Hz for the 𝐸 population and 14 Hz for the 𝐼 one as proposed in [28]), in our experiments we had to use higher set-point rates to attain reliable behavior, because of the relatively small network size and the low probability of connections used. In Table 1 we quantify the effects of network size and connectivity with additional experiments.

As the weights are shared globally within each population, the learning rules are implemented at the whole population level. Therefore the network maintains a given firing rate at the population level, rather than at the individual neuron level (see [28] for a local implementation of the rules). However, the heterogeneity in the analog circuits and their inherent noise produce diverse behaviors among neurons, for example leading some of the neurons to fire intermittently at slightly increased rates (while maintaining the desired set-point firing rate at the full population level). In the following section, we explain the training procedure on the hardware and how the changes in weight Δ𝑤 are translated to the fine 𝐹_𝑤_ and course 𝐶_𝑤_ synaptic bias parameters on chip.

We further extended our study and implemented the homeostatic rules proposed by Mackwood et al. [40]. The values for 𝑤_𝑒𝑒_ and 𝑤_𝑖𝑖_ remained unchanged during the learning process. We simplified the equations for the other two weights to ensure compatibility with the chip.

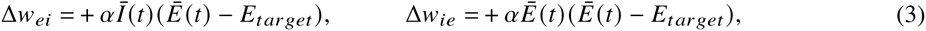

### Chip-in-the-loop training

Although the DYNAP-SE2 chip can be tuned to exhibit biologically plausible dynamics, it does not incorporate on-chip synaptic plasticity circuits that are capable of learning. To overcome this limitation we used a “chip-in-the-loop” learning paradigm: the activity produced by the chip is transmitted to a computer, which analyzes the data at the software level and calculates the parameter updates, which are then sent back to the chip.

In particular, the network is emulated for 1 s on the chip. Each emulation is repeated five times to reduce the influence of noise and obtain more reliable results. The output spikes are sent back to the computer, which averages the results, calculates the appropriate weight update, applies stochastic rounding (see below), and sends the updated weights to the chip for the next iteration (i.e., after 5 repetitions). After each iteration, we drain the accumulated residual current from both the neuron and synapse circuits. We stop the procedure after 400 iterations, as we found it to be sufficient for the network to converge to the target firing rates regardless of the initial conditions. Most simulations converge to target firing rate before iteration 200, which shows the long-term stability of the cross-homeostasis rules in a converged network.

For all experiments, we applied the same set of parameters to neurons and synapses at the beginning of the training, with the exception of the random initialization of weights. The training produced stable and reliable activity around the programmed set-points 100% of the time, without having to re-tune any of the neuron or synapse parameters (except the synaptic weights), across multiple days, and for multiple chips.

### Stochastic rounding

As the values of the synaptic weight parameters are set by the bias-generator DAC, the weight updates can only take discrete values Δ𝐹_𝑤_ = 0, ±1, ±2, For very large learning rates, this could lead to unstable learning, and for very low ones to no changes at all. To overcome the constraints of limited-resolution weight updates, a common strategy used is to use stochastic rounding (SR) [56, 57]. The SR technique consists of interpreting the Δ𝑤 calculated by the learning rule as a *probability* to increase or decrease the weight, as follows:

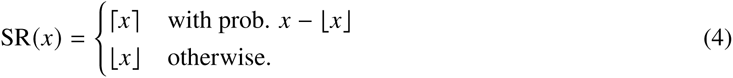

where ⌈·⌉ indicates the ceiling operation and ⌊·⌋ the floor operation.

Values for the desired weight updates Δ𝑤 are given by the learning rules (3). We also define upper and lower bounds for 𝐹_𝑤_, respectively 𝐹_−_ = 20 and 𝐹_+_ = 250. We then obtain the new coarse and fine values 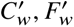 of the weight as follows:

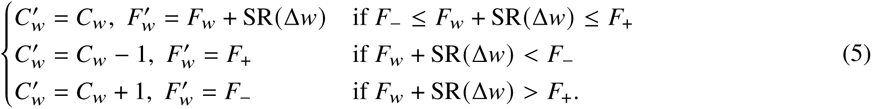

In other words, the fine value is updated with the stochastically-rounded version of Δ𝑤; when it oversteps the bound, the fine value is reset to the bound, and the coarse value is changed instead.

Note that this SR strategy is compatible with neuromorphic hardware implementations, as demonstrated in [58]. Our setup using a chip-in-the-loop enables the evaluation of different learning mechanisms and test the robustness of different design choices. The design of future on-chip learning circuit can therefore be informed by our results, following a hardware-software co-design approach.

## Author Contributions

Conceptualization: M, SSM and RL. Investigation: M, MS and SSM. Methodology, Software, Data curation and Visualization: M and MS. Supervision SSM, RL, DB and GI. Writing Original Draft: M and MS. Writing and editing: SSM, RL, DB and GI. Funding acquisition: SSM, MS, DB and GI.

## Acknowledgments

We thank the organizers of the CapoCaccia Workshop Towards Neuromorphic Intelligence 2022, where this work was conceived. We thank Chenxi Wu, Ole Richter and German Koestinger for DYNAP-SE2 support, and Emre 1. O. Neftci, Matthew Cook for useful discussions. SSM was supported by the Swiss National Science Foundation (SNSF) grant nos. P2ZHP3-187943 and P500PB-203133. M was supported by the SNSF Sinergia Project (CRSII5-180316). The work of MS was made possible by a postdoctoral fellowship from the ETH AI Center. GI was supported by the EU ERC Grant "NeuroAgents" (No. 724295), DVB and RL were supported by the National Institutes of Health grant no. NS116589. RL was supported by Universidad Nacional de Quilmes (Argentina), CONICET (Argentina), and NeurotechEU (Faculty Scholarship).

## Supplementary Material

**Table 3:** Pyramidal Neuron Specifications.

| Parameter | Coarse | Fine | Description |
| --- | --- | --- | --- |
| SOIF_LEAK_N | 2 | 4 | Membrane decay time constant |
| SOIF_GAIN_N | 3 | 40 | Neuron gain |
| SOIF_REFR_N | 2 | 45 | Refractory period |
| SOIF_DC_P | 0 | 0 | Constant DC current |
| SOIF_SPKTHR_P | 3 | 40 | Spike threshold |

**Table 4:** Connection Specifications for Pyramidal Neuron.

| Parameter | Coarse | Fine | Description |
| --- | --- | --- | --- |
| DEAM_ETAU_P | 2 | 30 | AMPA synaptic decay time constant |
| DEAM_EGAIN_P | 2 | 35 | AMPA synaptic gain |
| DEGA_ITAU_P | 1 | 50 | GABA-B (subtractive) decay time constant |
| DEGA_IGAIN_P | 1 | 50 | GABA-B gain |
| DENM_ETAU_P | 1 | 60 | NMDA decay time constant |
| DENM_EGAIN_P | 1 | 130 | NMDA gain |
| DESC_ITAU_P | 2 | 16 | GABA-A (shunt) decay time constant |
| DESC_IGAIN_P | 1 | 90 | GABA-A gain |
| SYPD_EXT_N | 3 | 200 | Pulse width |
| SYPD_DLY0_P | 0 | 0 | No Delay |
| SYPD_DLY1_P | 5 | 250 | Precise Delay |
| SYPD_DLY2_P | 1 | 80 | Mismatch Delay |

**Table 5:** PV Neuron Specifications, Parameter description is same as in 3.

| Parameter | Coarse | Fine |
| --- | --- | --- |
| SOIF_LEAK_N | 2 | 10 |
| SOIF_GAIN_N | 3 | 60 |
| SOIF_REFR_N | 2 | 85 |
| SOIF_DC_P | 0 | 0 |
| SOIF_SPKTHR_P | 3 | 60 |

**Table 6:** Connection Specifications for PV Neurons, Parameter description is same as in 4.

| Parameter | Coarse | Fine |
| --- | --- | --- |
| DEAM_ETAU_P | 2 | 30 |
| DEAM_EGAIN_P | 2 | 35 |
| DEGA_ITAU_P | 1 | 50 |
| DEGA_IGAIN_P | 1 | 70 |
| DENM_ETAU_P | 1 | 60 |
| DENM_EGAIN_P | 1 | 130 |
| DESC_ITAU_P | 2 | 16 |
| DESC_IGAIN_P | 1 | 90 |
| SYPD_EXT_N | 3 | 200 |
| SYPD_DLY0_P | 0 | 0 |
| SYPD_DLY1_P | 5 | 250 |
| SYPD_DLY2_P | 1 | 80 |

**Figure 9:**
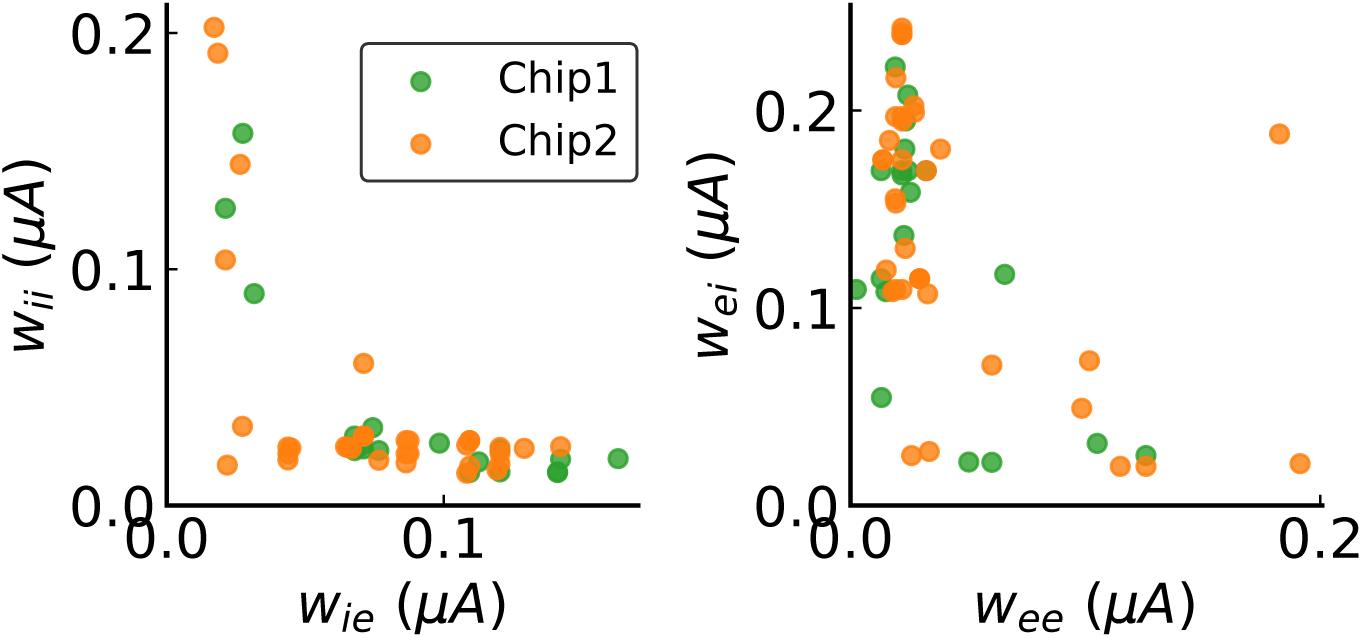
Initial weight configuration for experiments in Figure 2 e

**Figure 10:**
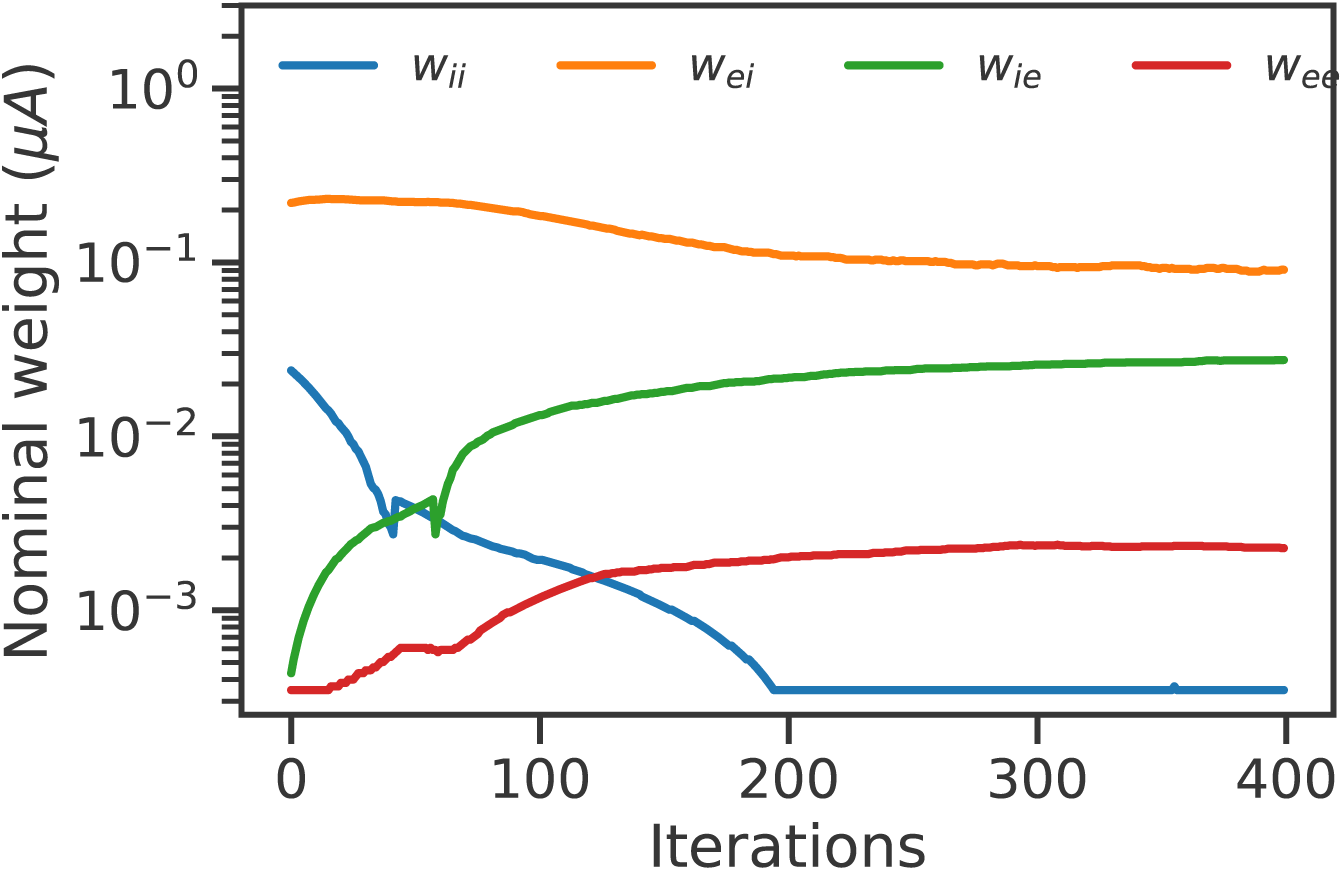
Another representative experiment (the same as in Figure 2) of the convergence of weight values. Here we show 𝑤_𝑒𝑒_ and 𝑤_𝑖𝑒_ were potentiated whereas 𝑤_𝑒𝑖_ and 𝑤_𝑖𝑖_ were depressed during learning.

**Figure 11:**
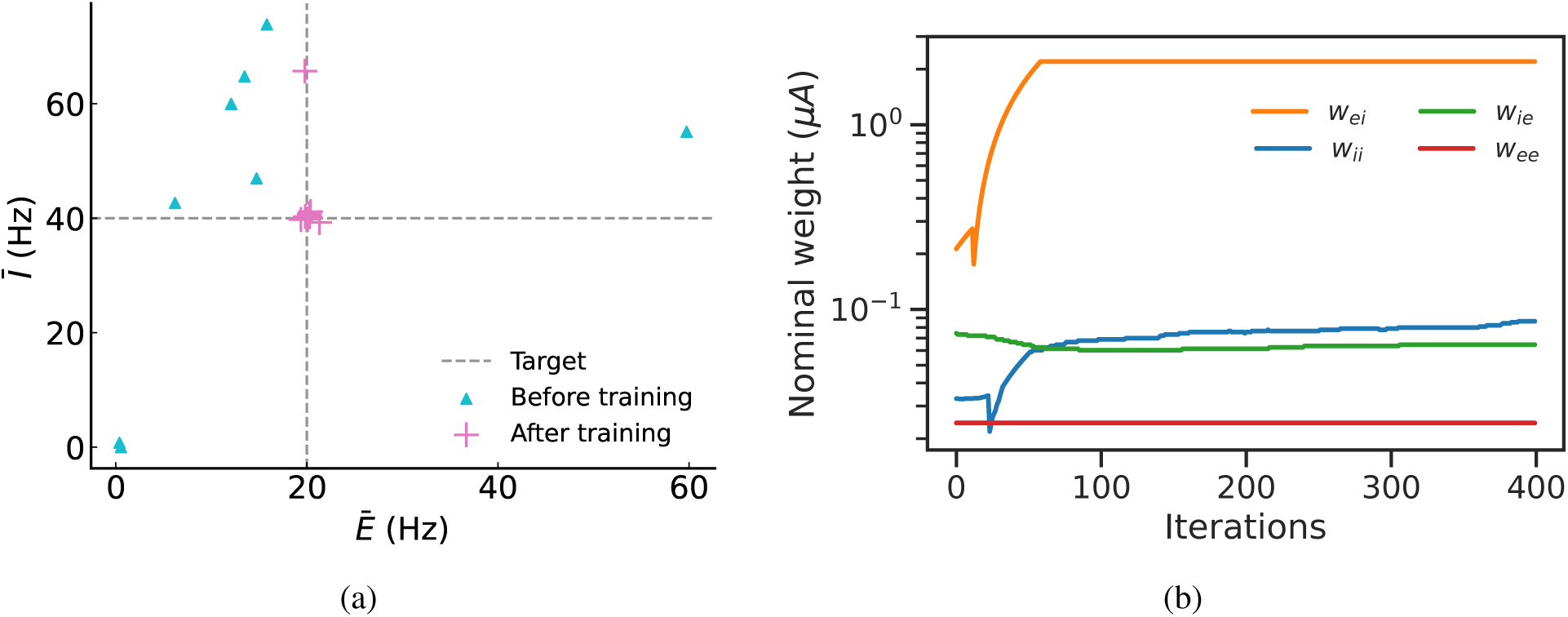
(a) Random networks ran with different initialization of all weights (n=8). Pyr → Pyr weights were froze during learning. Most firing rates converge to the targets, except in one of the experiments. (b) Weight dynamics during learning for the experiment that did not converge in (a). The 𝑤_𝑒𝑖_ weight reaches the maximum value permitted on chip, and the other weights are not able to compensate and fail to drive the firing rates to target.

**Figure 12:**
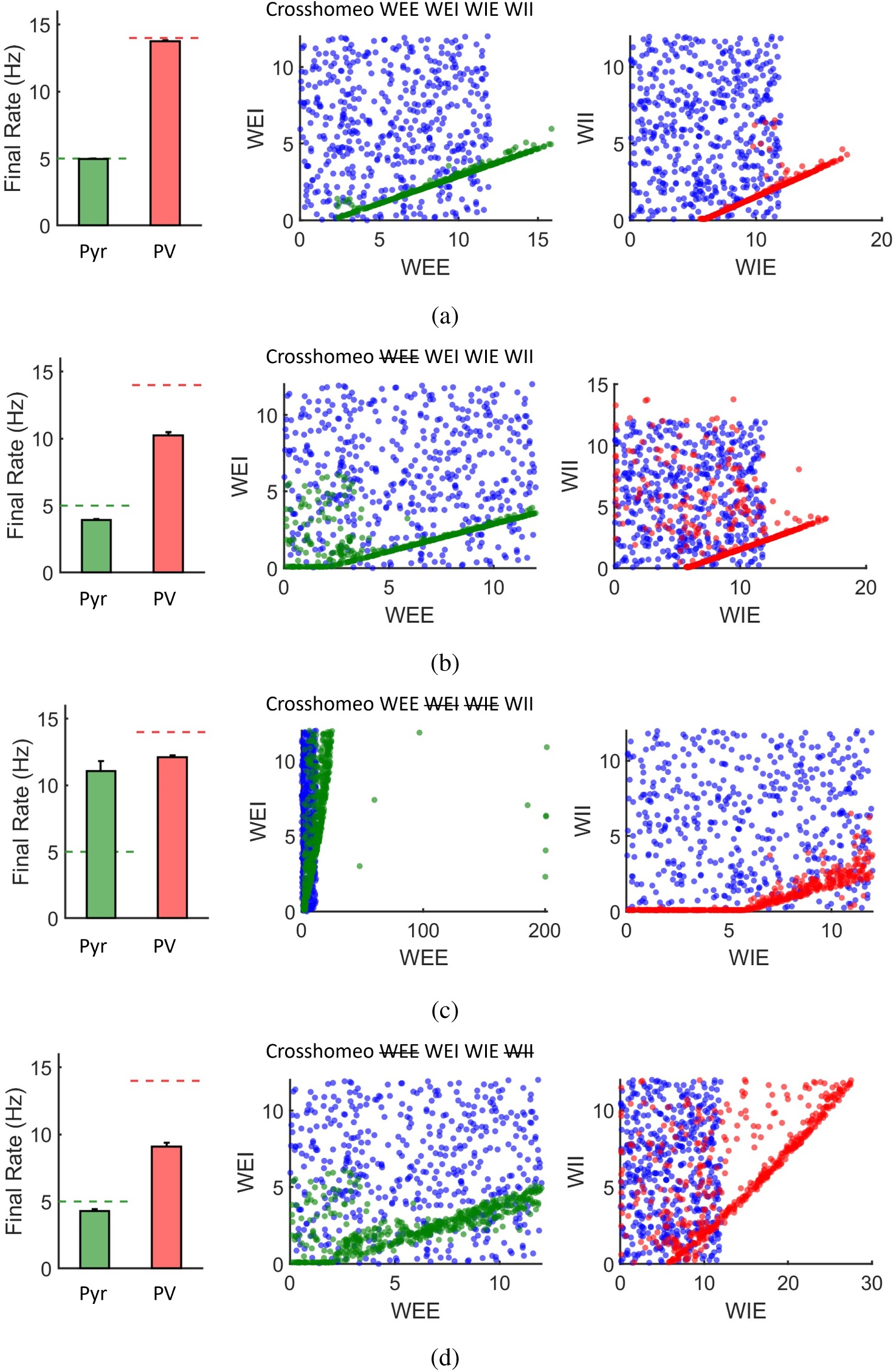
Simulations of a two-population E-I firing rate model under different combinations of cross-homeostatic rules. The rate model is described in [28]. Initial weights are randomly distributed in the interval [0, 12]. The column on the left shows final E and I rates (Pyr and PV) after 4500 trials, for n=500 experiments, with the dashed lines indicating target setpoints. The rest of the plots show weight values before (blue dots) and after (green and red dots) plasticity, over 4500 trials. Different combinations of cross-homestatic plasticity rules were tested: (a) Cross-homeostatic rules operating in the four weight classes 𝑤_𝑒𝑒_, 𝑤_𝑒𝑖_, 𝑤_𝑖𝑒_ and 𝑤_𝑖𝑖_ (b) Cross-homeostatic rules operating in the 𝑤_𝑒𝑖_, 𝑤_𝑖𝑒_ an 𝑤_𝑖𝑖_ weights (c) Cross-homeostatic rules operating in the 𝑤_𝑒𝑒_and 𝑤_𝑖𝑒_ weights (d) Cross-homeostatic rules operating in the 𝑤_𝑒𝑖_ and 𝑤_𝑖𝑒_ weights.

**Figure 13:**
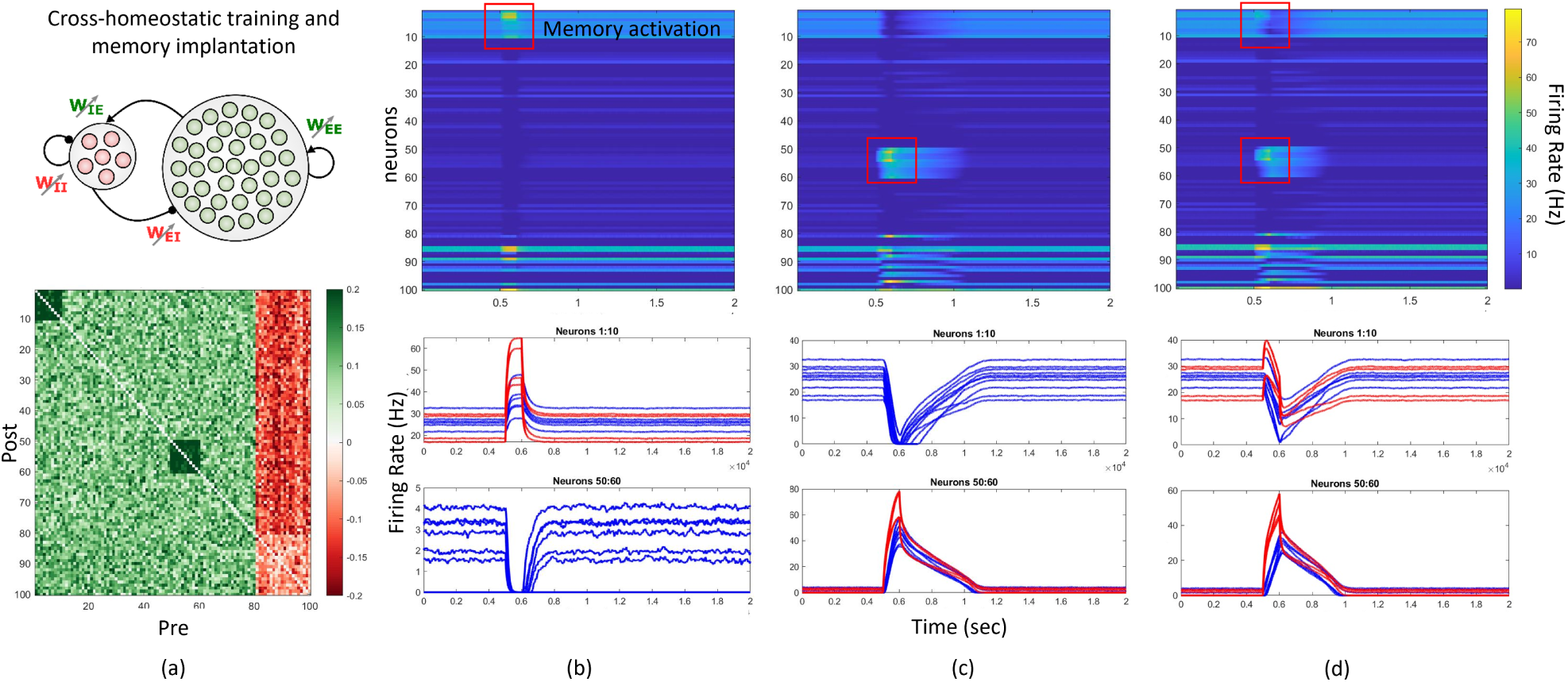
Cross-homeostasis implements lateral inhibition and sWTA dynamics in a firing rate network model. (a) Following [28] a firing rate network model of 80 excitatory and 20 inhibitory neurons is brought to stable self-sustained activity at its setpoints (𝐸_𝑠𝑒𝑡_ = 5 and 𝐼_𝑠𝑒𝑡_ = 14). After learning, two memories are implanted in the network, by artificially increasing the excitatory weights among a subset of excitatory neurons (neurons 1-10 and 50-60). (b) The implanted memory can be recalled by activating a subset of the neurons within the assembly (in red: neurons 1-4 for 100 ms). When recalling one of the memories, the second memory is effectively inhibited. Lateral inhibition is thus implemented by the cross-homeostatic rules. (c) The reverse is true when activating the second memory (in red: neurons 50-54). In this case, a form of working memory is triggered after the memory recall, as activity within the whole assembly lasts for 400 ms more after stimulus withdrawal. (d) When both memories are triggered (albeit with different stimulus intensity), the memory receiving the strongest input (in red: neurons 50-54) "wins", by effectively inhibiting the other assembly (neurons 1-10, neurons 1-4 in red receive input but of half the intensity of neurons 50-54). Cross-homeostasis therefore implements "for free", soft Winner Take All (sWTA) dynamics.

